# Contribution of Three tRNA Modification Enzymes to *Proteus mirabilis* Fitness and Catheter-Associated Urinary Tract Infection

**DOI:** 10.64898/2025.12.17.694933

**Authors:** Vitus Brix, Benjamin C. Hunt, Brian S. Learman, Aimee L. Brauer, Chelsie E. Armbruster

**Affiliations:** Department of Microbiology and Immunology, Jacobs School of Medicine and Biomedical Sciences, State University of New York at Buffalo, Buffalo, NY 14203, United States

## Abstract

RNA modifications play essential roles in cellular physiology by modulating RNA structure, stability, and translation efficiency. Among these, dihydrouridine is a conserved tRNA modification synthesized by the flavin-dependent enzymes DusA, DusB, and DusC, which introduces flexibility at defined uridine positions in the D-loop. While structural and biochemical studies in *Escherichia coli* have clarified Dus enzymes substrate specificities, their functional relevance in other bacterial pathogens remains uncharacterized. Here, we present the first functional characterization of the contribution of Dus enzymes to an important urinary tract pathogen, *Proteus mirabilis.* Insertional mutants were generated in *dusA, dusB*, and *dusC* in *P. mirabilis* strain HI4320 for evaluation of virulence-relevant phenotypes. All mutants grew similarly to wild-type under multiple culture conditions, although loss of *dusB* caused a fitness defect during growth in human urine. Disruption of *dusB* also resulted in increased biofilm biomass, impaired swimming motility, enhanced outer membrane permeability, and increased susceptibility to detergents and certain antibiotics, while disruption of *dusA* or *dusC* only impacted biofilm formation and susceptibility to detergents. In a murine model of catheter-associated urinary tract infection*, dusB* was also critical for fitness in all organs during co-challenge against wild-type as well as overall colonization and ascending infection during independent challenge. In contrast, *dusC* did not contribute to fitness and *dusA* showed a modest, compartment-specific contribution to fitness, as the *dusA::Kan* mutant was outcompeted in the bladder. Together, these findings identify DusB as a key regulator of motility, membrane integrity, and host fitness in *P. mirabilis*, linking tRNA modification to virulence.

## Introduction

Post-transcriptional RNA modifications are essential to RNA function, stability, and adaptability, serving as key regulators of gene expression and stress response across all domains of life(1, 2). In addition to their canonical roles in maintaining RNA structural integrity, emerging evidence highlights their involvement in modulating translation during environmental adaptation and virulence in pathogenic microbes(1, 3). Among these, tRNA modifications are the most chemically diverse, with over 100 different known base alterations that fine-tune tRNA folding, stability, and interactions with translation machinery(2, 4, 5). One such modification, dihydrouridine (D), is formed by the enzymatic reduction of uridine C5=C6 double bonds, which introduces flexibility by disrupting base stacking. This non-aromatic base modification is commonly found in the D-loop of tRNAs and promotes proper tertiary structure formation(1, 4).

Dihydrouridine is synthesized by dihydrouridine synthases (Dus), a family of flavin mononucleotide (FMN)-dependent enzymes conserved across bacteria, archaea, and eukaryotes (4, 6). In *Escherichia coli*, three Dus enzymes (DusA, DusB, and DusC) exhibit substrate-specific modification of uridine at positions 16, 17, and 20 of tRNAs, respectively, although full site specificity has only recently been defined (4, 7). Structurally, Dus enzymes share a conserved catalytic TIM barrel and a C-terminal helical domain but they differ in tRNA binding orientations, reflecting their substrate specificity(4, 8). Evolutionary analyses suggest that DusB is the ancestral enzyme, with DusA and DusC arising via gene duplication(4, 6).

Despite their conservation and structural characterization in model organisms, the functional roles of Dus enzymes in bacterial physiology, particularly in pathogenesis, remain largely unexplored. To date, no study has directly assessed Dus contributions in the context of any infectious disease model. However, there is increasing recognition that tRNA modifications influence bacterial virulence. For example, it has been shown that deletion of the tRNA prenyltransferase *miaA* significantly increases membrane permeability and antibiotic susceptibility in *E. coli* and reduces fitness in murine urinary tract infection models(9, 10). These findings highlight a broader role for tRNA-modifying enzymes in adaptation to host environments such as the urinary tract and underscore the need to examine other tRNA modifying enzymes in this context.

*Proteus mirabilis* is a Gram-negative, motile bacterium frequently implicated in complicated urinary tract infections (UTIs), and especially catheter-associated infections (CAUTIs)(11–13). The pathogenesis of *P. mirabilis* is driven by a combination of robust swarming motility, biofilm formation, resistance to host and environmental stressors, and urease activity (12, 13). These behaviors are tightly regulated at both the transcriptional and translational levels; for example, swarming requires precise control of flagellar biosynthesis, surface sensing, and stress response systems(14–16). Transcriptomic and proteomic studies have shown that different phases of swarming are marked by significant shifts in translation-related gene expression, including tRNA pools and ribosomal protein abundance(14, 15). Thus, *P. mirabilis* exhibits unique translationally-regulated behaviors and represents a powerful model to investigate how tRNA modifications contribute to virulence-associated phenotypes in uropathogens.

*P. mirabilis* strain HI4320 is a clinical isolate that encodes all three Dus enzymes (DusA, DusB, and DusC) at disparate chromosomal locations, and all three genes were previously identified by transposon insertion site sequencing as candidate fitness factors in a mouse model of CAUTI(17). DusB was also identified as a possible fitness factor for *P. mirabilis* ascending UTI in a signature-tagged mutagenesis study, and was upregulated during swarming compared to during growth in broth culture(16, 18). Here, we present the first functional characterization of the contribution of each Dus enzyme to *P. mirabilis* virulence-related phenotypes and fitness during infection. Using mutants of *dusA*, *dusB*, and *dusC*, along with plasmid-based complementation, we investigated the roles of the Dus enzymes in bacterial growth, biofilm formation, motility, membrane integrity, fitness, and ascending infection in the murine CAUTI model. Our findings reveal a non-redundant role for DusB in coordinating motility, maintaining outer membrane homeostasis, and supporting *P. mirabilis* fitness within the urinary tract while also contributing to colonization and ascension of the urinary tract during independent challenge. These results uncover novel links between RNA modification and virulence traits in *P. mirabilis* and suggest broader implications for Dus enzymes in pathogen adaptation to host environments.

## Methods

### Animal model ethics statement

Animal protocols were approved by the Institutional Animal Care and Use Committee of the University at Buffalo (MIC31107Y), in accordance with the Office of Laboratory Animal Welfare, the U.S. Department of Agriculture, and the Association for Assessment and Accreditation of Laboratory Animal Care. Mice were anesthetized with a weight-appropriate dose (0.1 ml for a mouse weighing 20 g) of ketamine/xylazine (80 to 120 mg/kg ketamine and 5 to 10 mg/kg xylazine) by intraperitoneal injection and euthanized by CO_2_ with vital organ removal.

### Bacterial strains

*Proteus mirabilis* strain HI4320 was originally isolated from the urine of a catheterized nursing home resident(19). All *dus* mutants were generated by inserting a kanamycin resistance cassette into the gene of interest using the Sigma TargeTron group II intron system according to the manufacturer’s instructions (Table 1)(19, 20). The *dusA*, *dusB*, and *dusC* mutants were verified by selection on LB agar containing kanamycin (50 µg/mL) and by colony PCR spanning the intron insertion site.

**Table 1.** Bacterial strains used in this study. Wild-type *Proteus mirabilis* HI4320, *dus* mutants, and corresponding complementation or empty vector strains are listed with their genetic modifications and antibiotic resistance profiles. Kanamycin resistance indicates successful targetron insertion, while ampicillin resistance corresponds to pGEN-based complementation or empty vector carriage.

| Strain Name | Description | Antibiotic Resistance |
| --- | --- | --- |
| ***Pm* HI4320** | Wild-type *P. mirabilis* | — |
| ***Pm* pGEN-EV** | Wild-type *Pm* with pGEN empty vector | Ampicillin |
| ***dusA*::Kan** | *dusA* disrupted by targetron insertion of kanamycin resistance cassette | Kanamycin |
| ***dusB*::Kan** | *dusB* disrupted by targetron insertion of kanamycin resistance cassette | Kanamycin |
| ***dusB* pGEN-EV** | *dusB* mutant with empty vector | Kanamycin, Ampicillin |
| ***dusB* pGEN-*dusB*** | *dusB* mutant complemented with *dusB* | Kanamycin, Ampicillin |
| ***dusC*::Kan** | *dusC* disrupted by targetron insertion of kanamycin resistance cassette | Kanamycin |

To control for potential polar effects caused by TargeTron retrohoming, each mutant was complemented by expressing the respective *dus* gene in trans from the native promoter(19). Complementation vectors were constructed by amplifying each full-length gene along with ∼500 bp of native upstream regulatory sequence and cloning into the multiple cloning site of the low-copy plasmid pGEN-MCS. Recombinant vectors were transformed into the respective mutant strains via electroporation and selected on LB agar containing ampicillin (100 µg/mL). The integrity of each construct was confirmed by colony PCR and Sanger sequencing.

### Culture conditions and media

*P. mirabilis* strains were routinely cultured at 37°C in low-salt LB (LSLB) broth (10 g/L tryptone, 5 g/L yeast extract, 0.1 g/L NaCl) with shaking at 225 rpm or on LSLB agar (1.5% agar). For selection, kanamycin (50 µg/mL) or ampicillin (100 µg/mL) were added as needed. Freezer stocks were prepared in 50% glycerol and stored at –80°C.

Artificial urine medium (AUM) was prepared to model the urinary tract environment *in vitro*(21). For additional physiologically relevant conditions, filtered pooled normal human urine was obtained from Lee Biosolutions (Catalogue number 991-03-P-FTD), aliquoted, and stored at -20°C. Prior to use, urine was thawed and diluted 1:1 with sterile 0.9% saline to reduce osmolarity and enhance reproducibility across batches. For defined minimal media experiments, *Proteus* minimal salts medium (PMSM) was prepared containing 10.5 g/L K_2_HPO_4_, 4.5 g/L KH_2_PO_4_, 1 g/L (NH_4_)_2_SO_4_, 15 g/L agar, supplemented with 0.002% nicotinic acid, 1 mM MgSO_4_, and 0.2% glycerol(19, 22).

### Growth Curve Assays

To assess *in vitro* growth kinetics, overnight cultures were centrifuged, washed with 1x PBS, and resuspended in LSLB, artificial urine medium (AUM), or diluted human urine at a starting inoculum of ∼1×10^7^ CFU/ml. For optical density monitoring of growth, 200 μL of each inoculum was pipetted into 96-well plates and OD_600_ readings were recorded every 15 minutes for 24 hours at 37°C with continuous orbital shaking using a SYNERGY-H1 plate reader. For determination of CFUs, 5 mL cultures were incubated at 37°C with shaking and samples were collected hourly, serially diluted, and plated using an Eddy Jet 2W Spiral Plater. CFU/ml was determined by counting colonies using a ProtoCOL 3 automatic zone-measuring colony counter.

### Co-challenge Growth Competition Assay

To assess relative fitness *in vitro*, co-challenge assays were performed by mixing wild-type (WT) and mutant strains at a 1:1 CFU ratio (∼5×10^6^ CFU/ml each) in 5 mL of fresh media. Cultures were incubated at 37°C with aeration for 7 hours. Samples were taken hourly, serially diluted, and plated on LSLB and LSLB with 50 µg/ml kanamycin to distinguish mutant and WT colonies. Competitive indices (CI) were calculated as the ratio of mutant to wild-type bacteria recovered from the output divided by the ratio of mutant to wild-type bacteria in the inoculum.

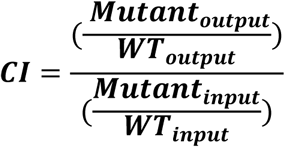

### Motility Assays

Swarming and swimming motility assays were performed using agar-based media prepared with defined nutrient concentrations and agar content to support flagellar-based movement. Swimming agar plates were prepared with 10 g/L tryptone, 5 g/L NaCl, and 2.5 g/L agar (0.25%) dissolved in deionized water and adjusted to pH ∼7. Swarming agar plates were prepared with 10 g/L tryptone, 5 g/L yeast extract, 5 g/L NaCl, and 13 g/L agar (1.3%), also dissolved in deionized water and adjusted to pH ∼7. Media were autoclaved and poured into 100 mm petri dishes under sterile conditions. Antibiotics were added as needed to maintain selection for complemented strains. Swim plates were stab-inoculated from overnight cultures and incubated at 30°C for ∼16 hours prior to measuring swimming diameter. Swarm agar plates were inoculated by spotting 5 µl of overnight culture on the center of plate, the inoculum was air-dried for 5-10 minutes, and swarm plates were incubated at 37°C for ∼16 hours prior to measuring swarm ring diameter.

### Biofilm Assays

Biofilm formation was quantified using crystal violet staining as previously described(23). Overnight cultures were diluted in the indicated medium and ∼1×10^7^ CFU/mL were seeded into 24-well plates (750 uL/well). After 24 hours at 37°C in humidified bags, supernatants were aspirated, wells were gently washed with 1×PBS, biofilms were fixed with 95% ethanol for 15 minutes and then air dried. Biofilms were then stained with 0.1% crystal violet for 1 hour, washed once with distilled water, remaining stain was solubilized in ethanol and wells were scraped with pipette tips. Biomass was quantified by absorbance measured at 570nm. For biofilm CFU enumeration, biofilms were instead washed once with PBS, fully suspended in 1 mL PBS by scraping the sides and bottom of the well, serially diluted, and plated.

### Membrane Permeability Assays

Bacterial membrane integrity was assessed via uptake of the hydrophobic fluorescent dye N-phenyl-1-naphthylamine (NPN)(19). Overnight cultures were washed and resuspended in HEPES buffer (5 mM HEPES, 5 mM glucose, pH 7.2). Conditions included buffer only (BUF+NPN), buffer with bacteria (BAC+NPN), and buffer with bacteria and 1 mM EDTA (BAC+EDTA+NPN). Bacterial suspensions were adjusted to OD_600_ ∼0.5, and 10 uM NPN was added. Fluorescence (Ex/Em: 350/420 nm) was measured using a SYNERGY-H1 plate reader. Each assay was performed in triplicate, and the NPN uptake factor was calculated as follows:

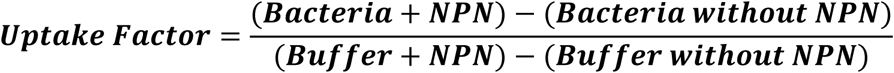

### Antibiotic and Detergent Susceptibility

Disk diffusion assays were performed to assess the sensitivity of *P. mirabilis* wild-type and *dus* mutant strains to detergents and antibiotics. Overnight cultures were adjusted to an OD600 of 0.5 and spread uniformly on LSLB agar plates using sterile swabs. Sterile 6 mm paper disks were pre-impregnated with 10 uL of sodium dodecyl sulfate (SDS) or sodium deoxycholate (DOC) at 2.5%, 5%, and 10% concentrations and allowed to dry briefly before being placed on the agar surface. Antibiotic susceptibility was assessed using pre-loaded commercial disks containing polymyxin B (300 units), ceftazidime (30 ug), imipenem (10 ug), ciprofloxacin (5 ug), and cefoxitin (30 ug). Plates were incubated at 37°C for 24 hours, and inhibition zone diameters were measured in millimeters using digital calipers.

### Murine CAUTI Infection Model

To evaluate the *in vivo* fitness of *P. mirabilis dus* mutants, 6-to 8-week-old female CBA/J mice were transurethrally inoculated with a total of 1×10⁵ CFU in 50 μL of either a single strain (independent challenge) or a 1:1 mixture of WT and mutant strains (co-challenge). A sterile 4-mm silicone catheter segment was inserted into the bladder during inoculation to simulate catheter-associated urinary tract infection (CAUTI) conditions. At either 48 or 96 hours post-infection, mice were euthanized and bacterial burdens were quantified from urine, bladder, kidneys, and spleens. Organs were aseptically harvested, weighed, and mechanically homogenized in sterile PBS using a Bullet Blender 5 Gold tissue homogenizer (Next Advance). Homogenates and urine were serially diluted and plated on LSLB agar using an EddyJet 2 spiral plater (Neutec Group). Selective antibiotics (kanamycin/ampicillin) were included when necessary to differentiate mutant and wild-type in co-challenge experiments. Plates were incubated at 37 °C for 16–18 hours before colony enumeration using a ProtoCOL 3 automated colony counter (Synbiosis). Competitive indices were calculated as above.

### Statistical Analysis

Statistical analysis was performed using GraphPad Prism version 10.5 (GraphPad Software). For biofilm quantification, disc diffusion, membrane permeability, growth curves, and CFU data, significance was determined using two-way analysis of variance (ANOVA) with appropriate corrections for multiple comparisons. Competitive index values from co-challenge experiments were analyzed using the Wilcoxon signed-rank test. All experiments were conducted with at least three biological replicates unless otherwise noted. A *p*-value of <0.05 was considered statistically significant.

## Results

### DusB Contributes to *P. mirabilis* Fitness *In Vitro*

To assess whether any of the dihydrouridine synthases contribute to *P. mirabilis* growth, we first compared the *in vitro* growth kinetics of *P. mirabilis* HI4320 wild-type (WT), *dusA::Kan, dusB::Kan,* and *dusC::Kan* across multiple media types (Figure 1A-C). In low salt LB (LSLB, rich medium), *Proteus* minimal salts medium (PMSM), and artificial urine medium (AUM), all strains displayed similar lag times, exponential growth rates, and final cell densities over 18 hours, indicating that none of the *dus* genes are important for growth under standard culture conditions. Likewise, no growth differences were observed in pooled human urine over a 7-hour time course (Figure 1D), confirming that none of the *dus* genes are critical for *P. mirabilis* proliferation under physiologically relevant conditions *in vitro*.

**Figure 1.**
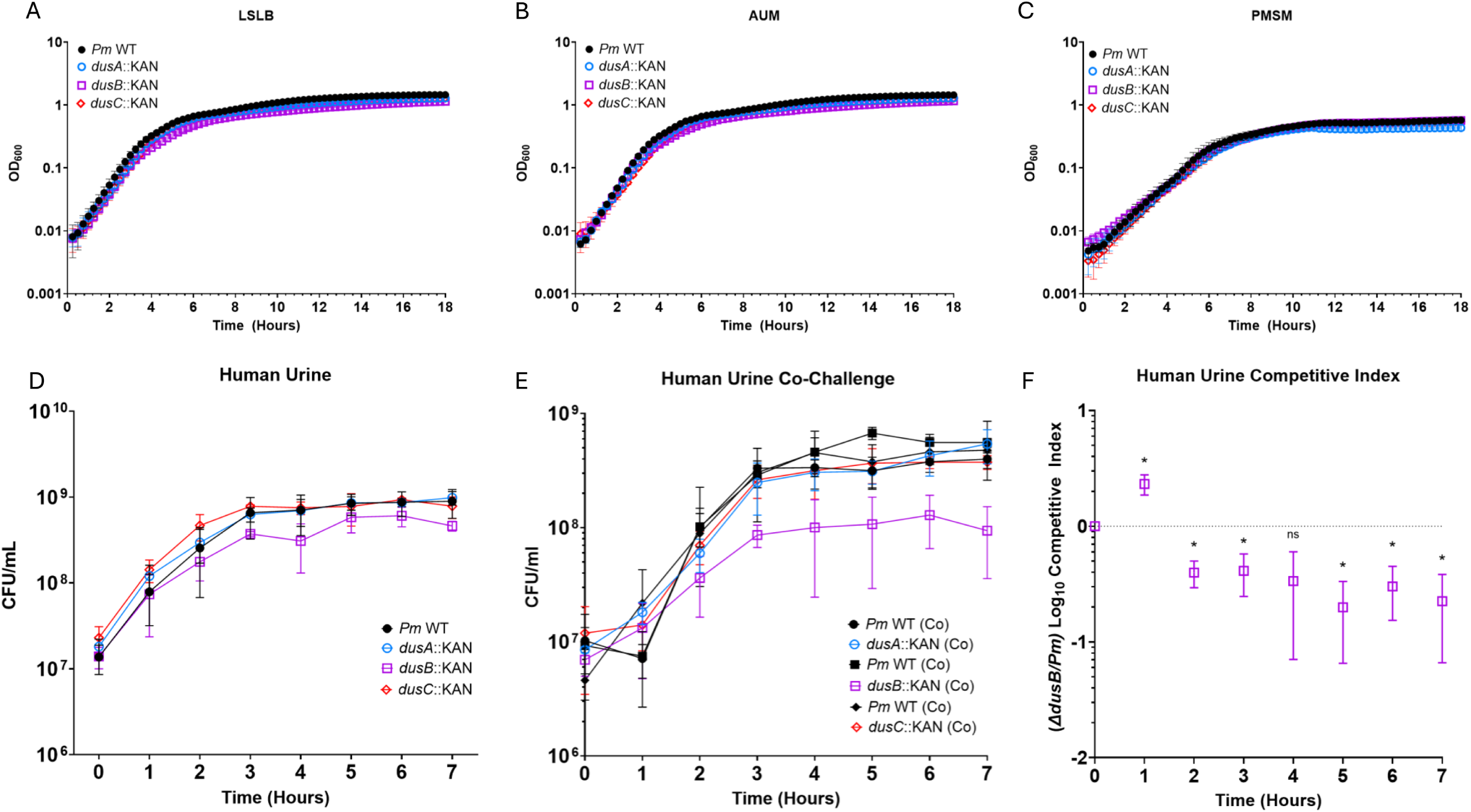
Loss of *dusB* causes a competitive growth defect in co-culture but grows normally in isolation. **(A–C)** Growth of wild-type *Proteus mirabilis* (WT), *dusA::Kan*, *dusB::Kan*, and *dusC::Kan* was assessed by optical density over 18 hours in low salt LB (LSLB), artificial urine medium (AUM), or *Proteus* minimal salts medium (PMSM). **(D)** Growth of wild-type and mutant strains in pooled human urine was assessed by plating for CFUs at hourly time points over 7 hours. **(E)** Fitness defects of the *dus* mutants were assessed by inoculating human urine with a 1:1 mixture of wild-type and the indicated mutant, and CFUs were assessed by differential plating at hourly time points over 7 hours. **(F)** Competitive indices (CI) for *dusB::Kan* were calculated from the same *in vitro* urine co-challenge assay shown in **(E)**, revealing a CI < 1 consistent with decreased fitness during co-challenge. The dotted line marks a CI of 1 (equal fitness). Data are shown as mean ± SD with at least 3 biological replicates performed for each; significance was assessed via two-way ANOVA with multiple comparison correction (panels A–E) or one-sample t-test (panel F) using GraphPad Prism v10.5.

To next probe whether any of the dihydrouridine synthases contribute to *P. mirabilis* fitness under host-relevant conditions stress, we performed co-culture competition assays in human urine (Figure 1E). Notably, the *dusB::Kan* mutant was significantly outcompeted by WT starting at ∼2 hours post-inoculation, while *dusA::Kan* and *dusC::Kan* showed no competitive disadvantage (Figure 1F). These findings reveal that DusB is dispensable for individual growth but contributes to bacterial fitness when resources are limited or when competing strains are present.

### Dus Mutants Display Altered Biofilm Formation Across Media Conditions

Biofilms are a critical contributor to *P. mirabilis* colonization, persistence, and pathogenesis in the catheterized urinary tract, providing protection from host immune responses and increasing resistance to antimicrobial agents(13, 24, 25). To determine whether dihydrouridine synthase mutants exhibit altered biofilm phenotypes, we assessed biofilm biomass and viability across multiple media conditions. In LSLB, all three mutants exhibited a robust increase in crystal violet staining relative to WT, with *dusB::Kan* showing slightly lower levels than the other two yet still significantly elevated compared to WT (Figure 2A). Despite having enhanced biomass, overall viability of the biofilm-associated bacteria was similar between *dusA::Kan, dusC::Kan,* and WT while viability of *dusB::Kan* exhibited a slight though statistically-significantly reduction (Figure 2D). Biofilms established in artificial urine medium (AUM) recapitulated the LSLB trend, including the decrease in *dusB::Kan* CFUs, although viability of *dusA::Kan* and *dusC::Kan* was slightly increased compared to WT (Figures 2B & 2E). In human urine, biofilm biomass of the *dusA::Kan* and *dusC::Kan* mutants exhibited a modest though statistically significantly increase compared to WT while *dusB::Kan* was indistinguishable from WT (Figure 2C). Viability measurements mirrored this trend, with *dusA::Kan* and *dusC::Kan* biofilms having elevated CFUs and *dusB::Kan* being identical to WT (Figure 2F). Taken together, these results demonstrate that loss of any of the dihydrouridine synthases generally increases biofilm accumulation to some degree, that increased biofilm biomass typically corresponds to increased CFUs when *dusA* or *dusC* are disrupted but not when *dusB* is disrupted. Thus, tRNA modifications by the dihydrouridine synthases may play unique roles in coordinating matrix production and cell survival during biofilm formation.

**Figure 2.**
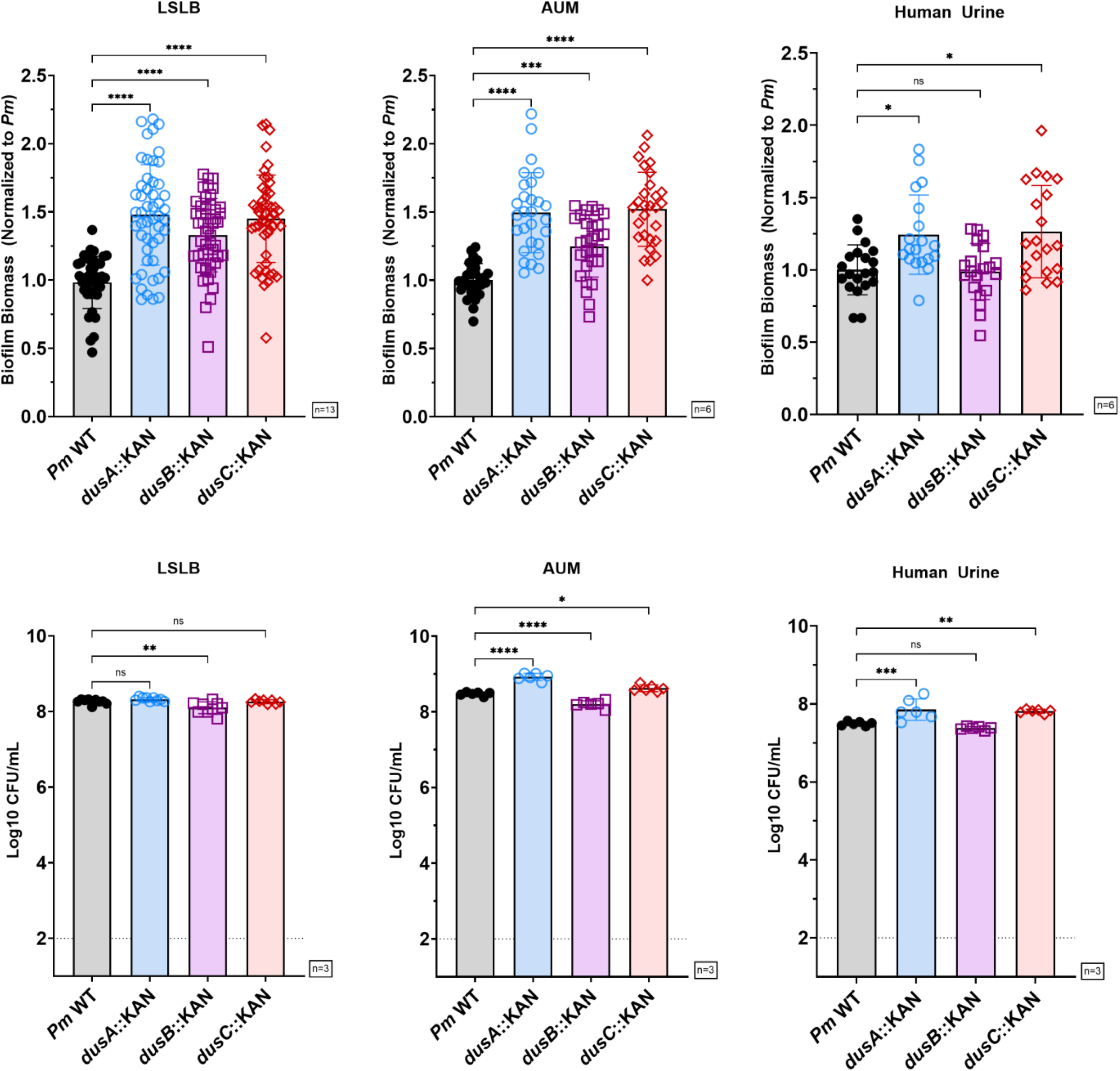
Biofilm biomass and viability of *P. mirabilis dus* mutants across media conditions. (**A–C**) Biofilm biomass was quantified by crystal violet staining after 20 hours of static growth in LSLB (**A**), artificial urine medium (AUM) (**B**), and pooled human urine (**C**). (**D–F**) Viable counts of biofilm-associated bacteria from the same conditions as panels A–C were determined by serial dilution and plating to assess CFU in LSLB (**D**), AUM (**E**), and pooled human urine (**F**). Each point represents a biological replicate; bars indicate the median. Statistical significance was assessed by Kruskal-Wallis test with post-hoc Dunn’s multiple comparisons.

### DusB Selectively Impacts Swimming Motility Without Affecting Swarming Motility

To assess the impact of dihydrouridine synthase disruption on motility in *P. mirabilis*, we evaluated both swimming and swarming behaviors in wild-type (WT) and *dus* mutants. In soft agar swim assays, the *dusB::Kan* mutant exhibited a marked reduction in swimming halo diameter relative to WT and the other mutants, a phenotype that was restored upon complementation with pGEN-*dusB*, confirming specificity (Figures 3A, Supplemental Figure S1). Neither *dusA::Kan* nor *dusC::Kan* displayed defects in swimming motility compared to WT (Figures 3A, Supplemental Figure S1). Swarming motility, measured by the number and spacing of concentric rings formed on hard agar, revealed no significant alterations in overall swarm morphology. All strains consistently produced four concentric rings with the outermost reaching the plate edge, although *dusB::Kan* displayed a modest reduction in the diameter of the first swarm ring that could indicate a potential delay in swarm initiation (Figures 3B, Supplemental Figure S2). Together, these data demonstrate that DusB plays a selective and specific role in regulating swimming motility, while swarming behavior remains largely unaffected.

**Figure 3.**
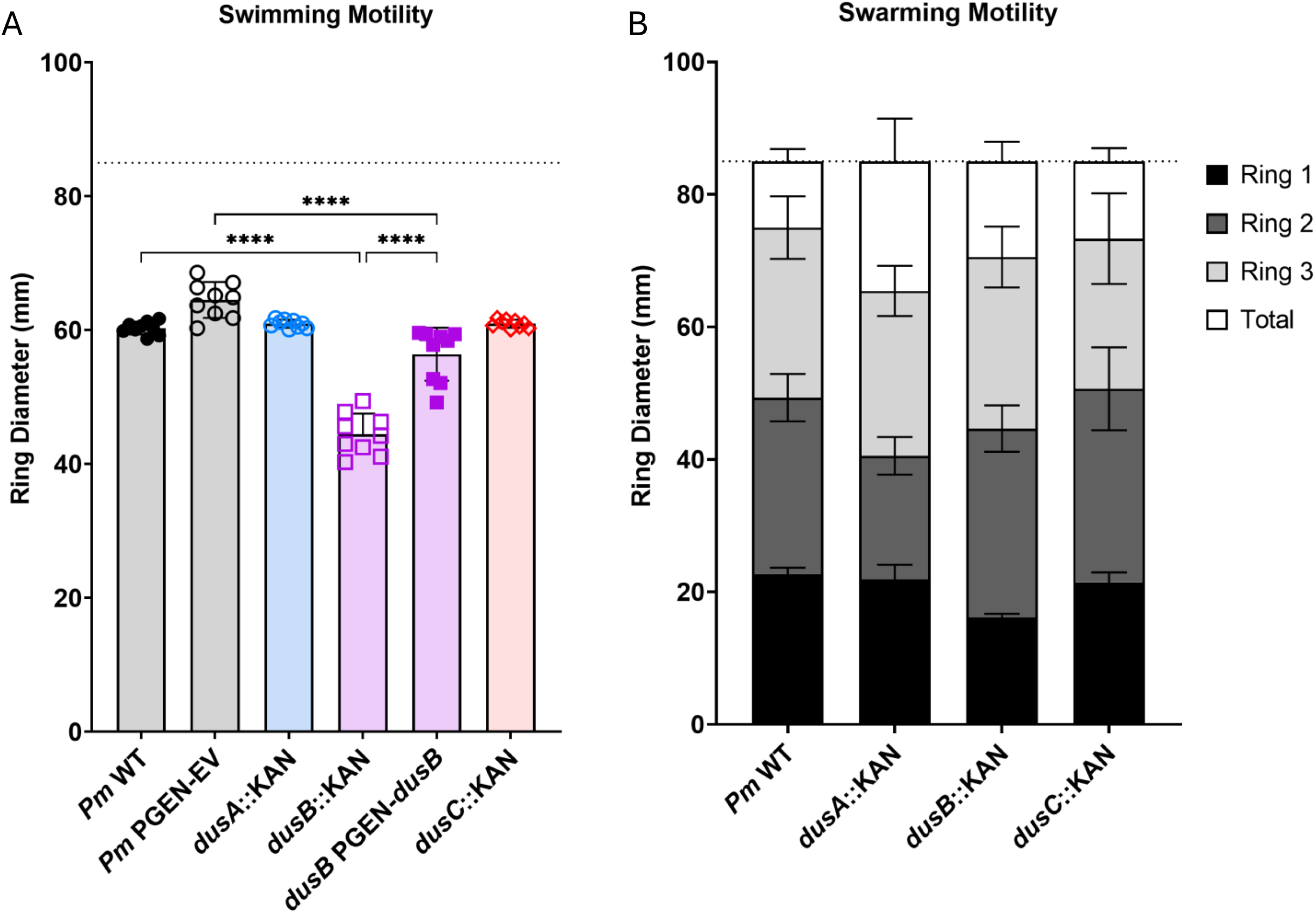
Assessment of Swimming and Swarming Motility in *dus* Mutants and Complementation Strains. (**A**) Swimming motility was quantified by measuring swim ring diameter on 0.25% agar plates after 18 hours of incubation. (**B**) Swarming motility was assessed on 1.5% agar swarm plates by measuring the diameter of four concentric swarm rings. Data represent mean ± SD from three independent experiments. Statistical analysis was performed using one-way ANOVA with multiple comparisons.

### Disruption of DusB Increases Outer Membrane Permeability

Motility defects can arise from disruption of the proton motive force (PMF), which is essential for flagellar rotation and is directly influenced by membrane potential and permeability(26, 27). As such, altered outer membrane integrity can compromise energy homeostasis and bacterial motility, making it a relevant phenotype to assess alongside swarming and swimming behavior(26–29). To determine whether *dus* gene disruption affects membrane integrity, we performed 1-N-phenylnaphthylamine (NPN) uptake assays (Figure 4). WT *P. mirabilis* exhibited low NPN uptake under basal conditions, consistent with an intact outer membrane that excludes hydrophobic probes, and a significant increase upon exposure to EDTA. The *dusB::Kan* mutant displayed elevated permeability that was restored to WT levels upon complementation. Neither *dusA::Kan* nor *dusC::Kan* showed any difference from WT under these conditions. These results identify DusB as a key contributor to outer membrane stability.

**Figure 4.**
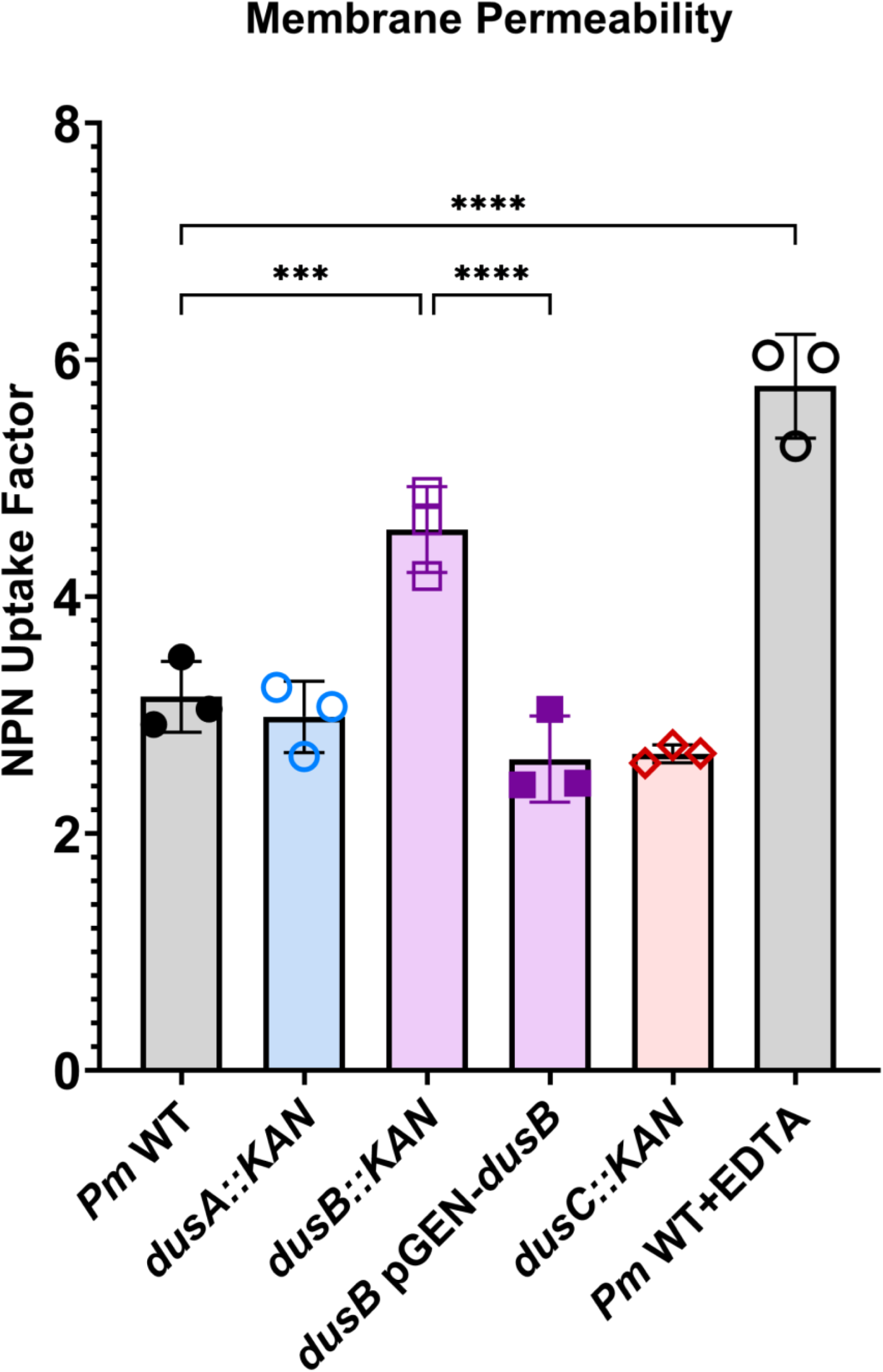
NPN uptake assays reveal increased membrane permeability in *dusB::Kan*. Outer membrane integrity of *P. mirabilis* WT and *dus* mutants was assessed using 1-N-phenylnaphthylamine (NPN) uptake. Fluorescence intensity was measured for cells incubated with NPN alone (buffer + bacteria + NPN) or following EDTA treatment (buffer + bacteria + EDTA + NPN), which destabilizes the outer membrane by chelating divalent cations. Error bars represent mean ± SD from three independent experiments. Statistical significance was determined using one-way ANOVA.

### Dus Mutants Exhibit Increased Sensitivity to Membrane-Disrupting Detergents

To determine whether *dus* deletion affects outer membrane integrity, we assessed the susceptibility of each mutant to sodium dodecyl sulfate (SDS) and sodium deoxycholate (DOC), both of which are detergents known to disrupt membrane integrity. Zones of inhibition were measured for wild-type *P. mirabilis* and the *dus* mutants on LSLB plates with discs impregnated with various concentrations of detergents (Figure 5A). WT was not inhibited by 2.5% SDS or by any concentration of DOC, but a zone of inhibition was preset at 5% and 10% SDS. All of the *dus* mutants exhibited zones of inhibition at 2.5% SDS, indicate that all were more susceptible to membrane disruption: however, the *dusB::Kan* mutant consistently exhibited the largest zone of inhibition and was significantly enlarged compared to WT at 5% and 10% SDS, indicating increased sensitivity to membrane disruption. For DOC, none of the mutants were sensitive at 2.5% but all displayed zones of inhibition at 5% and 10%. Again, *dusB::Kan* showed the largest zones of inhibition, highlighting its heightened vulnerability to detergent-mediated stress. These findings indicate that all three Dus enzymes contribute to maintaining outer membrane integrity, with DusB playing the most prominent role. Complementation experiments were not performed due to concerns with maintaining antibiotic selection of plasmids in the presence of membrane-disrupting detergents.

**Figure 5.**
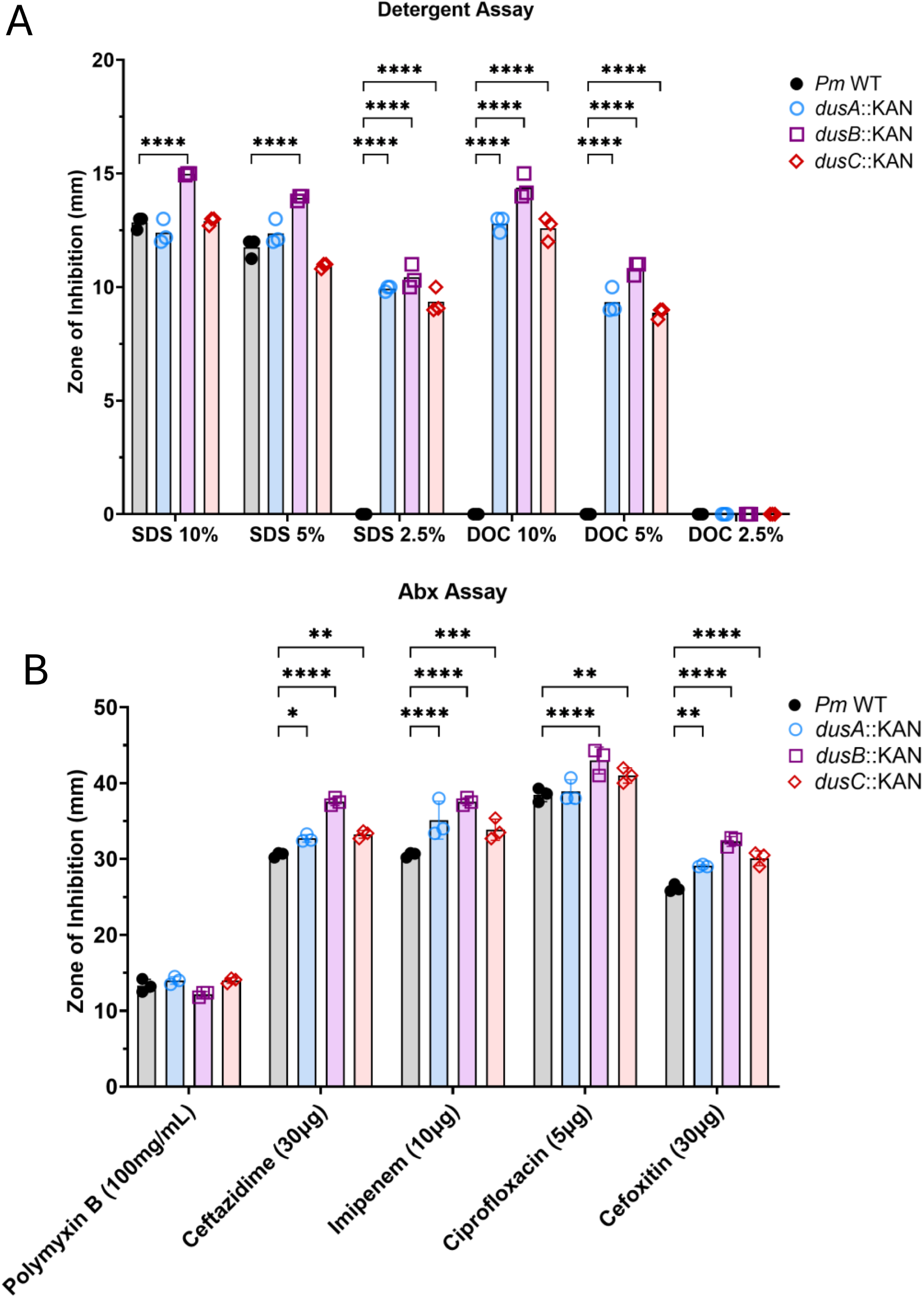
Sensitivity of *P. mirabilis dus* mutants to antibiotics and detergents. (A) Detergent sensitivity was tested by placing discs impregnated with SDS or sodium deoxycholate (DOC) solutions onto agar plates that were swab-inoculated to generate a bacterial lawn. Zones of inhibition were measured after overnight incubation. (B) Antibiotic susceptibility was evaluated by disc diffusion using standard antibiotic-impregnated discs. Data shown are from three independent experiments. Statistical significance was determined by one-way ANOVA with multiple comparisons.

Next, we assessed susceptibility to antibiotics with differing mechanisms of action (Figure 5B). The *dusB::Kan* mutant showed significantly enlarged zones of inhibition for cell wall synthesis inhibitors ceftazidime, cefoxitin, and imipenem, and the DNA replication inhibitor ciprofloxacin. *P. mirabilis* is intrinsically resistant to membrane-targeting cationic peptide antibiotics in the polymyxin class and there was no change in susceptibility of the *dus* mutants to polymyxin B, suggesting that the observed susceptibility pattern is not due to generalized outer membrane destabilization. The *dusA::Kan* and *dusC::Kan* mutants showed more modest changes in antibiotic susceptibility than *dusB::Kan*; both were more susceptible than wild-type to the same antibiotics as *dusB::Kan,* although *dusA::Kan* did not exhibit altered sensitivity to ciprofloxacin. Together, these findings suggest that the tRNA modifications catalyzed by DusB are important for maintaining outer membrane integrity and resistance to both membrane-disrupting agents and antibiotics with intracellular targets, while the modifications catalyzed by DusA and DusC have a lesser contribution.

### DusB Contributes to Fitness and Ascending Infection in the Catheterized Urinary Tract

To evaluate whether *dus* enzymes influence the ability of *P. mirabilis* to colonize and compete within a host environment, we employed a murine model of CAUTI(11). Female CBAJ mice were transurethrally implanted with 4mm silicone catheter segments and co-inoculated with 1:1 mixtures of WT and *dus* mutant strains. After 96 hours, bacterial burden was quantified from urine, bladder, kidneys, and spleen to calculate competitive indices (CI) (Figure 6). The majority of mice co-challenged with *dusA::Kan* showed a moderate reduction in *dusA::Kan* CFUs in the urine and bladder compared to WT, but the competitive index only demonstrated a statistically significant fitness defect in the bladder (Figure 6A). In contrast, the *dusB::Kan* mutant was significantly outcompeted by WT in all four compartments and was below the limit of detection in roughly half of all mice, indicating a pronounced *in vivo* fitness defect (Figure 6B). This experiment was repeated with an endpoint of 48 hours to determine the temporal contribution of *dusB* to *P. mirabilis* fitness, and *dusB::Kan* was again recovered at the limit of detection in roughly half of the mice and exhibited a significant defect in all organs (Supplemental Figure 3). The *dusC::Kan* mutant displayed no significant competitive defect in any compartment (Figure 6C). These findings demonstrate that DusB is essential for fitness of *P. mirabilis* in the CAUTI model even at earlier time points post inoculation, likely due to its combined roles in membrane integrity, motility, and contribution to fitness during growth in urine. The compartment-specific defect observed for *dusA::Kan* suggests a more localized contribution to bladder colonization, whereas *dusC::Kan* appears dispensable under these infection conditions.

**Figure 6.**
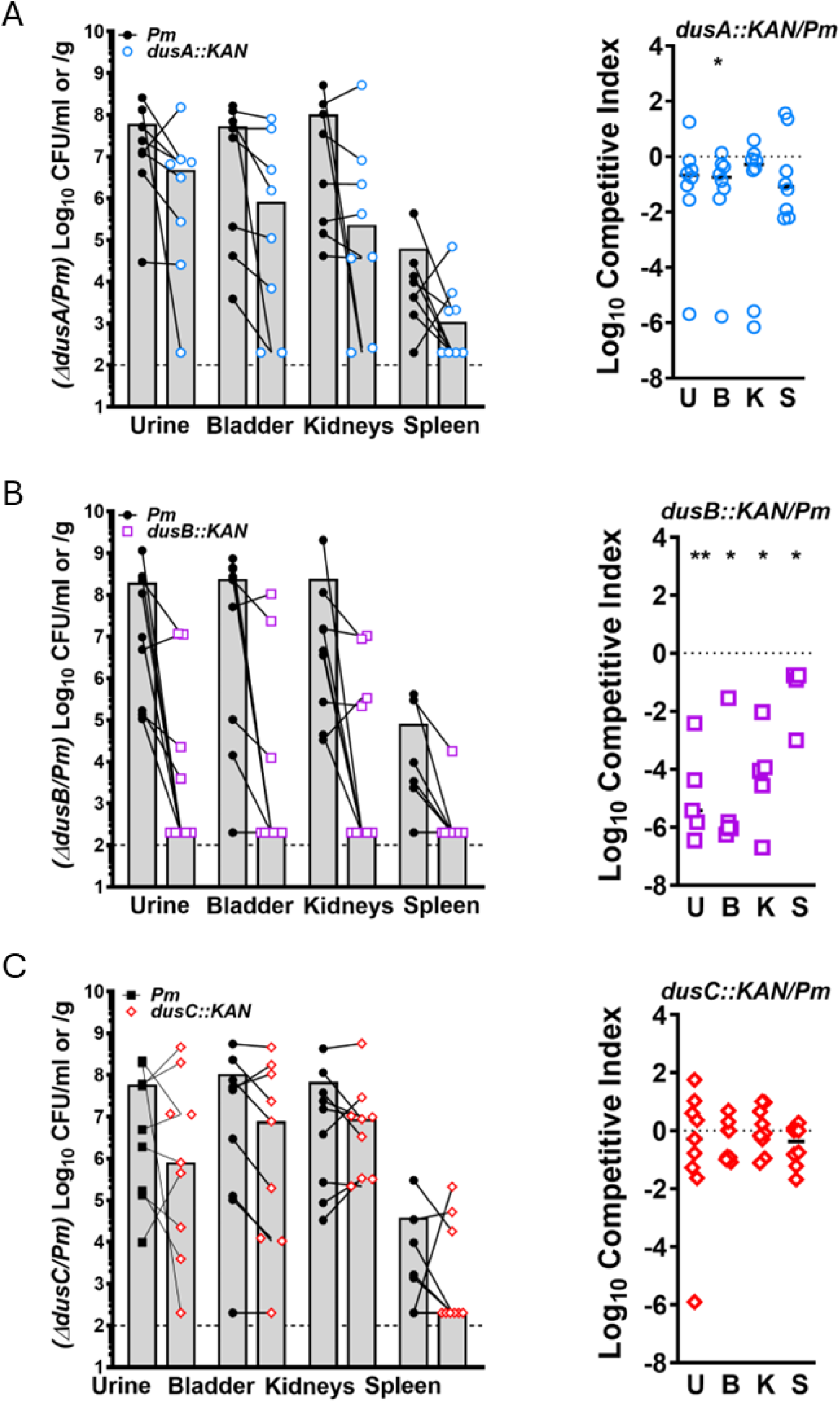
Dus enzymes contribute differentially to *P. mirabilis* fitness during murine CAUTI. Female CBA/J mice were transurethrally catheterized and co-infected with 1x10^5^ CFUs of a 1:1 mix of WT *P. mirabilis* and the indicated *dus* mutant (A: *dusA::Kan*, B: *dusB::Kan*, C: *dusC::Kan*). Bacterial burdens were enumerated from urine, bladder, kidneys, and spleen after 96 h. Left panels show CFU/g or CFU/ml for WT and mutant strains from the same mouse connected by a black line. Right panels show calculated competitive indices (mutant/WT). Statistical significance was determined by Wilcoxon signed-rank test.

We next sought to determine whether DusB is critical for *P. mirabilis* colonization and ascending infection or if it only contributes to fitness during competition with another strain. Female CBA/J mice were catheterized and inoculated with either wild-type *P. mirabilis* or *dusB::KAN,* and bacterial burden was assessed in the urine, bladder, kidneys, and spleen 96 hours post-infection (Figure 7). During independent challenge, mice inoculated with *dusB*::KAN exhibited a significant reduction in bacterial burden in the urine, bladder, and kidneys compared to mice inoculated with wild-type *P. mirabilis,* indicating that DusB is an important contributor to colonization of the urinary tract and ascension to the kidneys in the CAUTI model. In contrast, bacterial burdens in the spleen were comparable between strains, indicating that disruption of *dusB* does not significantly impair systemic dissemination from the urinary tract despite being required for fitness within the urinary tract and spleen during co-challenge.

**Figure 7.**
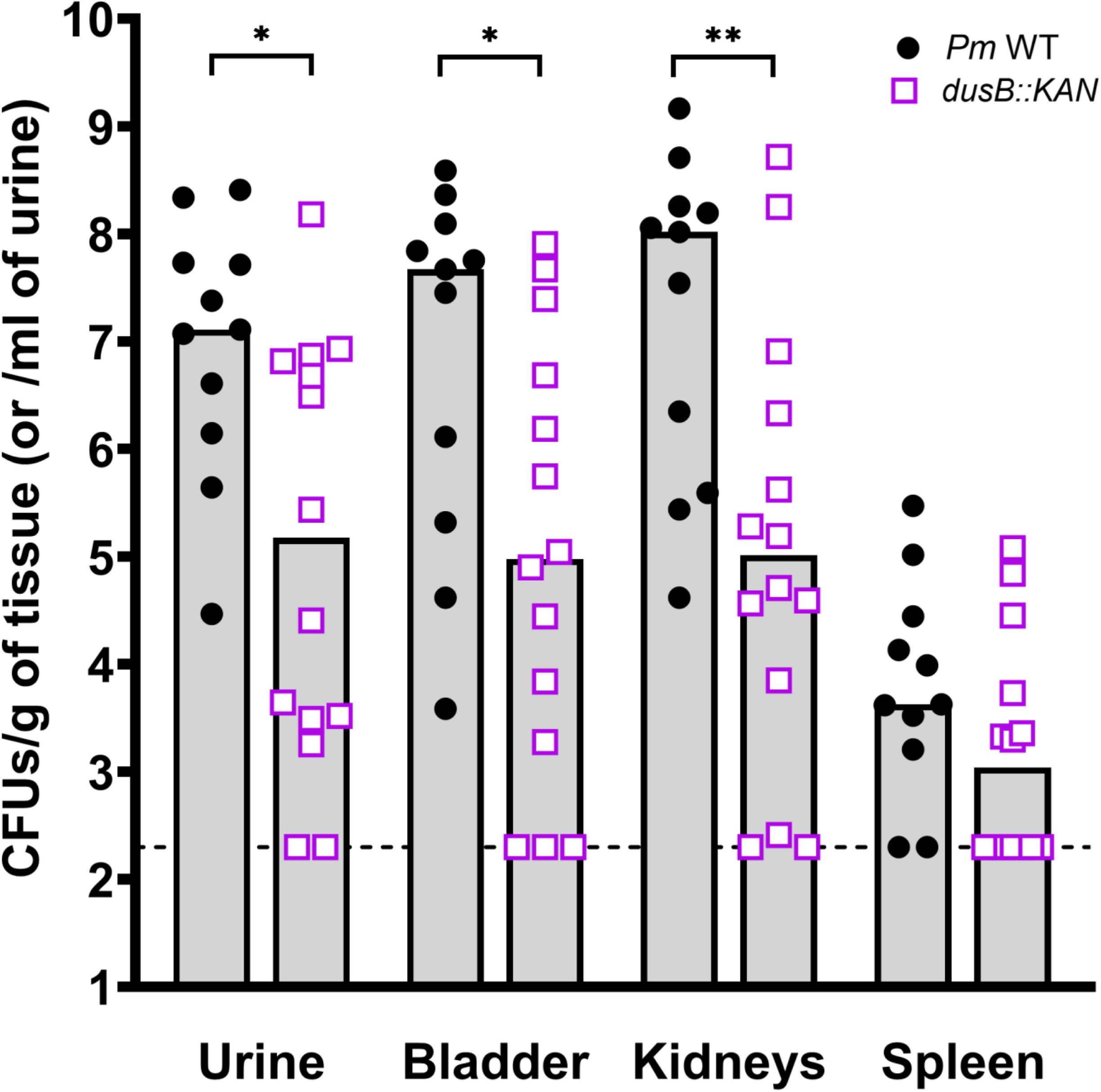
DusB contributes to *P. mirabilis* colonization and ascending infection during independent challenge. CBA/J mice were transurethrally catheterized and inoculated with 1×10⁵ CFU of either *P. mirabilis* HI4320 wild-type (WT) or *dusB::KAN*. After 96 hours, urine was collected, and organs were harvested, weighed, homogenized, and spiral plated for bacterial enumeration. Data points represent individual mice; bars show median. Statistically significance assessed via One-Way ANOVA.

## Discussion

This study presents the first investigation of dihydrouridine synthase (Dus) enzymes in the uropathogen *Proteus mirabilis*, revealing distinct and non-redundant roles for *dusA::Kan*, *dusB::Kan*, and *dusC::Kan* in modulating physiology and pathogenesis-associated traits. While Dus enzymes have been structurally and biochemically studied in model organisms such as *Escherichia coli*(7, 8), their functional roles in bacterial virulence remain largely unexplored. Our findings highlight the importance of DusB in regulating biofilm formation, motility, membrane stability, fitness, and ascending infection, with more subtle contributions from DusA and DusC.

Previous work in *E. coli* identified the substrate specificities of DusA, DusB, and DusC in modifying uridines at positions 20, 17, and 16 of the D-loop, respectively(4, 7). The incorporation of dihydrouridine increases tRNA flexibility, potentially modulating translation dynamics under different environmental stresses(1). In our study, loss of *dus* genes did not alter *in-vitro* growth kinetics even in minimal media, consistent with earlier studies where *dus* deletions had limited effects on basal viability in laboratory conditions(7, 30). However, we did uncover a fitness contribution of *dusB* to growth in the host-like environment of human urine, which may demand efficient translation of stress response factors to cope with nutrient limitation and altered osmolarity. The loss of *dusB*-mediated tRNA modification could impair fine-tuning of adaptive responses to these stressors, leading to decreased fitness under physiologically relevant conditions.

All *dus* mutants exhibited increased crystal violet staining, indicating elevated biofilm formation under static conditions. However, this increase did not correspond to a uniform rise in viable cell counts. Specifically, *dusA::Kan* and *dusC::Kan* mutants showed increased CFU recovery from the biofilm, while the *dusB::Kan* mutant displayed decreased CFUs despite elevated biomass. This discrepancy suggests that biofilm matrix components or dead cells may contribute to biofilm biomass in the *dusB* mutant, whereas the *dusA* and *dusC* mutants may have increased biomass due to enhanced adherence or reduced dispersal of the biofilm. These distinct outcomes highlight potential differences in how each Dus enzyme modulates survival versus surface association and raise the possibility that altered tRNA modifications influence differential partitioning between planktonic and adherent populations. On possibility is that Dus-mediated tRNA modifications may differentially fine-tune the translation of proteins involved in biofilm regulation, including stress response regulators or surface adhesins. Similar associations between translation efficiency and adaptation to host-like environments have been observed in other pathogens, underscoring the broader role of translational control in modulating virulence traits(3). For example, in *Pseudomonas aeruginosa*, tRNA modifications such as queuosine and m¹G impact the translation of stress response and quorum sensing genes, which are directly tied to biofilm development and antibiotic tolerance(31). Dus-mediated modulation of tRNA structure in *P. mirabilis* could similarly influence translation of biofilm-associated regulators, although the exact targets remain to be defined.

A striking finding of this study was the selective motility phenotype of the *dusB::Kan* mutant, which displayed a clear defect in swimming motility but largely preserved swarming behavior. Swimming and swarming motility in *P. mirabilis* both depend on tightly regulated expression and function of flagella, with swarming also involving regulatory pathways involved in differentiation into elongated swarm cells(32, 33). Altered tRNA modifications in *dusB::Kan* may therefore impair translation of specific swimming-associated factors while leaving flagellar biosynthesis and the broader swarming regulator program intact. Additionally, the swimming defect may be compounded by the increased membrane permeability observed in *dusB::Kan*, since intact outer membranes are essential for stable flagellar motor anchoring and for maintaining proton motive force(34). Together, these results indicate that DusB contributes specifically to swimming motility, while swarming is largely unaffected.

Increased sensitivity of *dusB::Kan* to detergents and antibiotics further supports its involvement in cell membrane or envelope integrity. We observed that *dusB::Kan* was more susceptible to SDS, DOC, and multiple antibiotics targeting the cell wall, but not to polymyxin B. This suggests that DusB-dependent pathways may influence outer membrane composition or porin expression, but not overall membrane charge or the lipid A modifications targeted by polymyxin. NPN uptake assays confirmed that *dusB::Kan* has increased baseline outer membrane permeability. These findings support the emerging concept that tRNA-modifying enzymes can impact membrane integrity and antibiotic susceptibility(35).

Critically, the *in vivo* CAUTI model revealed that *dusB* is required for fitness in all tested compartments (urine, bladder, kidneys, and spleen) demonstrating its essential contribution to survival during co-challenge. This finding is consistent with the previous genome-wide mutagenesis screens that identified *dusB* as a candidate fitness determinant in UTI and CAUTI models, further validating its *in vivo* relevance(17). During independent challenge, the *dusB*::KAN mutant also exhibited significantly reduced bacterial burdens in urine, bladder, and kidneys, indicating a clear role in colonization and ascending infection in addition to a role in fitness. Importantly, this attenuation was not observed in the spleen during independent challenge, suggesting that loss of *dusB* does not broadly impair systemic survival or dissemination, but instead selectively compromises colonization of niches encountered within the urinary tract.

The CAUTI environment imposes multiple stressors, including osmotic shifts, nutrient limitation, and host immune responses including exposure to antimicrobial peptides(36, 37)(12, 17, 33). If the tRNA modifications carried out by DusB are critical for rapid translation of proteins involved in responding to these stressors, loss of *dusB* could severely impair the ability of *P. mirabilis* to adapt to these stressors. The modifications carried out by DusB appear to be the most critical for *P. mirabilis* fitness, as disruption of either DusA or DusC has a minimal impact on *in vivo* fitness and less pronounced impacts on *in vitro* phenotypes. The compartment-specific attenuation of *dusA::Kan* may also reflect differential requirements for dihydrouridine-modified tRNAs in supporting stress responses or adherence-related pathways within the bladder. It is possible that DusA, DusB, and DusC each contribute to modification of distinct subsets of tRNAs, thereby influencing separate regulatory or structural protein pools required in a compartment-specific manner. If so, this would align with evolutionary data showing that DusB is likely the ancestral enzyme from which *dusA::Kan* and *dusC::Kan* arose via gene duplication(6). The stronger phenotypes observed with *dusB::Kan* support its central and possibly broader role in core tRNA modification, while *dusA::Kan* and *dusC::Kan* may have evolved more specialized functions.

Together, this work demonstrates that Dus enzymes, particularly DusB, are important regulators of biofilm formation, motility, membrane integrity, and *in-vivo* pathogenesis in *P. mirabilis*. By linking dihydrouridine synthesis to bacterial fitness, we reveal a novel layer of post-transcriptional control that shapes virulence traits in a clinically relevant pathogen. Future work should investigate the specific translational targets of Dus-modified tRNAs under infection-relevant conditions using approaches such as proteomics, ribosome profiling or tRNA modification sequencing. Additionally, understanding the intersection of tRNA modifications with stress response pathways may uncover new strategies for targeting RNA-modifying enzymes in antimicrobial development.

## Acknowledgments

This work was funded by the National Institute of Diabetes and Digestive and Kidney Diseases under award R01 DK123158 to CEA. The content is solely the responsibility of the authors and does not necessarily represent the official views of the National Institutes of Health.

## Author contributions

V.B, B.C.H., and C.E.A. designed experiments, analyzed data, and prepared the manuscript. V.B., B.C.H., B.S.L., and A.L.B. performed experiments. All authors reviewed the manuscript.

